# Host species and body site equally impact microbiome structure between sympatric Atlantic sea cucumber species

**DOI:** 10.64898/2025.12.12.694001

**Authors:** Catherine A. Crowley, Bernard S. Belisle, Emily Dart, Nathan A. Ahlgren

**Affiliations:** Clark University, Biology Department, Worcester, Massachusetts, USA; University of Connecticut, Department of Marine Sciences, Groton, Connecticut, USA

## Abstract

Sea cucumbers are key members in the marine ecosystem, food webs, and support valuable commercial fisheries, yet their microbiomes, which likely influence host health and function, are relatively understudied. There is specifically a paucity of microbiome studies of echinoderm coelomic fluid (CF), which is centrally involved in circulation, digestion, and immunity. This study analyzed the microbiomes of two sympatric holothurians (sea cucumbers), *Leptosynapta tenuis,* and *Sclerodactyla briareus*, found in coastal, temperate waters of the Northeast Atlantic. First, we found unexpectedly high levels (approaching 10^7^ cells/ml) of prokaryotic-sized cells in the coelomic fluid of the sympatric sea star *Asterias forbesi*. Amplicon sequencing of the 16S rRNA gene revealed significant differences in alpha and beta diversity of sea cucumber microbiomes. This included differences in CF communities between the two sea cucumber species and among body sites—CF, epidermis, and gut—of *S. briareus*. Host and body sites explained roughly equal amounts of microbial community variation (22% and 27%, respectively). We identified particular taxa associated with differences in community composition, including enrichment of anaerobic taxa (e.g., Desulftobacterota) in gut samples and Proteobacteria (including Burkholderiales and Pseudomonadales) in epidermis samples. We also uncovered higher levels of Synechococcales cyanobacteria in CF relative to the gut, epidermis, and background seawater samples. CF is thought to exchange with surrounding seawater, so these results highlight an apparent selective retention of CF microbial communities. This work adds to our growing knowledge of echinoderm microbiomes and highlights the need for additional surveys of their microbiomes.

**Importance:** Microbiomes influence the function and health of animal hosts, but basic characterization of sea cucumber microbiomes is lacking. Furthermore, there are few studies on the microbiomes of echinoderm coelomic fluid that interacts with the respiratory, digestion, locomotion, and reproduction systems of these animals. To fill these gaps in knowledge, we analyzed the microbiomes of two sea cucumber and one sea star species from the North Atlantic. We found unexpectedly high abundances of bacteria-sized cells in the coelomic fluid of all three species. Using DNA sequencing, we uncovered differences in bacterial communities across the two sea cucumber species and across body sites (the epidermis, gut, and coelomic fluid) within one host species. This is consistent with patterns of host and body-site specificity seen in animal other microbiome studies. These results build important information for sea cucumbers that play important roles in natural marine ecosystems and support globally important fisheries.

## Introduction

Echinoderms are predominantly ocean-floor-dwelling invertebrates that are classified by their five-fold radial symmetry, water vascular system, and spiny skin (1). The phylum Echinodermata is divided into five classes, including Asteroidea (sea stars), Echinoidea (sea urchins and sand dollars), Holothuroidea (sea cucumbers), Ophiuroidea (brittle stars), and Crinoidea (sea lilies and feather stars) (1). Echinodermata play a key role in ecosystem health and are vital to the marine food web and carbon cycles (2–5). Echinoderms use a variety of feeding strategies that contribute to their success in a wide range of environments. These include filter-feeding, grazing, hunting, and the digestion of detritus (6).

Echinoderms have societal importance as resources for both fisheries and research. Many echinoderms are harvested or cultured for human consumption (4, 7); sea cucumbers alone support US$1.4 billion annual trade worldwide (8). They have also been used as model organisms in biological, medical, and genetic research, (9, 10) including for their ability to regenerate parts of their body (11, 12). Echinoderm microbiomes have been relatively understudied, but important work is emerging on the impacts of microbes on echinoderm development, organ regeneration, and aquaculture (13–15).

Microbes form associations with every animal on the planet, and these associations play key roles in the ecology, physiology, and health of animals (16). Microbiome research has primarily focused on the influence of microbes found in the digestive tracts, and these microbes impact organismal health and function (17). This is in part because these sites are primary entry points for infection and because microbes play key roles in digestion (17). Distinct aspects of echinoderm anatomy provide a unique system to study host-microbe interactions. Echinoderms have a skeleton made of interlocking calcium carbonate plates and spines that are integrated with mucus-producing epidermal cells and provide unique substrates for microbial colonization (1).

Echinoderms possess a multifunctional coelom, a tissue-lined cavity filled with coelomic fluid (CF) (1, 18, 19). The coelom provides internal structure to the organism, drives locomotion via the water vascular system; transports nutrients (20) and respiratory gases (21); and supports the immune system (22). Similar to the blood in humans, the CF shuttles nutrients and immunocytes throughout the body, but unlike blood, CF contains high levels of microbes (> 10^5^ cells/ml) (19, 23, 24). The CF microbiome therefore likely plays important roles in the health and function of echinoderms.

There are a relatively limited number of studies on holothurian microbiomes (13, 25–34), and most are focused on gut microbiomes of commercially important sea cucumbers, e.g., *Apostichopus japonicus*, *Holothuria scabra,* and *Isostichopus badionotus*. There are only two reports of holothurian CF microbial communities (26, 34), despite the important connections CF has to multiple systems in these organisms. Several studies have shown that echinoderm communities differ across body sites (e.g., CF, skin, gut) and distinctly from surrounding seawater (19, 26, 35, 36). Different host species commonly harbor distinct microbiomes, but there are only a few comparative studies across echinoderm species (19, 36). These studies do find host-associated differences in microbiomes, but studies often only compare two species (36). A notable exception is Jackson et al. (2018), which surveyed twelve Pacific sea star species. We also note there are only a few studies of native temperate Atlantic echinoderms (13, 33); most prior studies are on tropical and/or Pacific species. Furthermore, there are limited reports on the abundance of bacteria in the CF of holothurians or echinoderms more broadly (19, 23, 24).

To help fill in these gaps in knowledge of echinoderm microbiomes, we characterized prokaryotic microbiomes across three body sites—epidermis, gut, and CF—of three sympatric species found in the Northwest Atlantic: one sea star species, *Asterias forbesi*, and two sea cucumber species, *Leptosynapta tenuis* and *Sclerodactyla briareus*. The two sea cucumber species are strict filter feeders, while the sea star species can actively hunt, filter-feed, and degrade detritus. This study sought to assess possible differences in microbiomes across body sites, between host species, and between organisms and surrounding seawater. We report unexpectedly high levels of bacteria in the CF of these species. We also find differences in CF microbiomes among the two sympatric sea cucumber species.

## Results

### Microbial cell abundance in coelomic fluid

Coelomic fluid (CF) was collected from six individuals within three sympatric echinoderm species found in coastal waters of the Northeastern Atlantic: *Leptosynapta tenuis, Sclerodactyla briareus, and Asterias forbesi*. The first two are sea cucumber species (class Holothuroidea) and the latter is a sea star species (class Asteroidea). Background seawater samples were also collected with each species. Significant differences in prokaryotic-sized cell abundances among CF and seawater samples were detected from ANOVA (F=5.8, =0.013) and subsequent Tukey’s test results. Background seawater had a mean cell abundance of 1.6×10^6^, which is typical for surface seawater. CF abundances in *L. tenuis* and *S. briareus* were not significantly different from background seawater or each other (mean: 1.5 x10^6^ and 1.1 x10^6^, respectively; p>0.05, Tukey’s test). CF abundances in *A. forbesi* however were significantly higher than the other CF samples and background seawater (p<0.05, Tukey’s test) with a mean of 7.5×10^6^ or roughly five-fold higher (Fig. 1).

**Figure 1.**
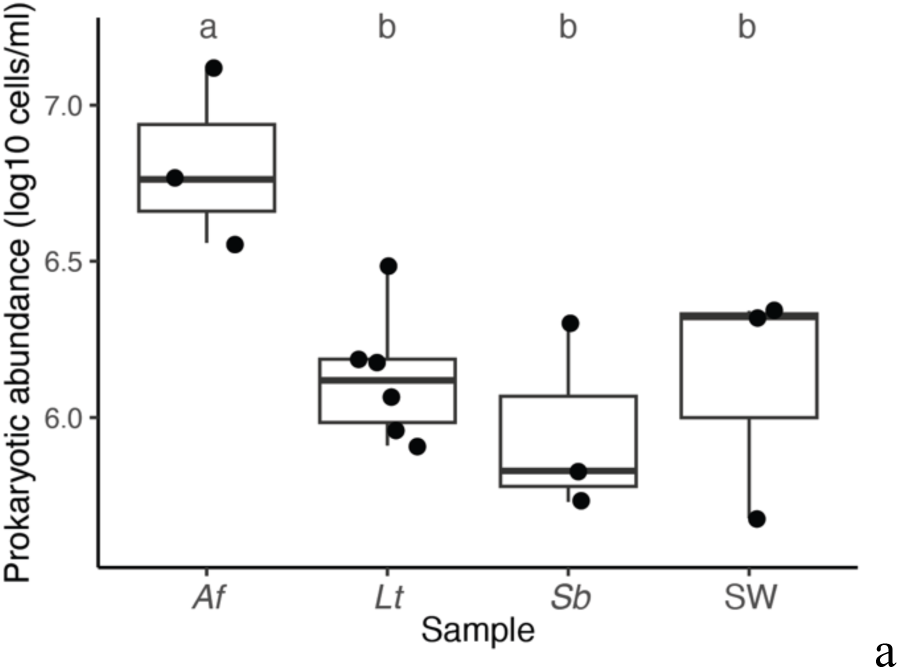
Prokaryotic cell abundances in the CF of echinoderms as determined by flow cytometry with Sybr Green I-staining. Horizontal lines indicate median values, boxes show interquartile ranges, and whiskers are 1.5 times the interquartile ranges. Sample abbreviations: *Af* = *Asterias forbesi*, *Lt* = *Leptosynapta tenuis*, *Sb* = *Sclerodactyla briareus*, SW = background seawater sample. Letters indicate if mean values were significantly different from each other (p<0.05, Tukey’s test) when the letters differ between sample categories.

### Microbial community alpha diversity patterns

Out of 42 microbiome samples collected from three body sites (epidermis, CF, and gut) across the three echinoderm species, only 22 generated sufficient 16S rRNA amplicon reads for community analysis (≥ 8000 reads). Unfortunately, no *A. forbesi* (Af) samples exceeded this threshold. There were CF samples from four *L. tenuis* (Lt) individuals, but only a single usable epidermis sample. For *S. briareus* (Sb), sufficient reads were generated for several individuals across the three sites: epidermis (*n*=5), gut (*n*=5), and CF (*n*=6). These outcomes limited the statistical comparisons that could be done among Lt samples and across the two host species, but there were sufficient samples to compare CF communities between Lt and Sb.

Rarefaction analysis shows plateauing of curves for all samples, supporting that sequencing had sufficiently sampled the majority of diversity present (Fig. 2A). A total of 13,886 ASVs were detected across all samples with 1175 ± 859 ASVs per sample (mean ± s.d.; range: 84-3031). Comparing CF samples, Lt had higher Shannon diversity than Sb (p=0.016, t-test) (Fig. 2B). For Sb samples, ANOVA detected significant differences between sites (F=14.0, p=0.006). Epidermis sites had significantly lower Shannon diversity values than CF and gut samples (p<0.005, Tukey’s test), but there was no significant difference in Shannon index values between the latter two sites (p=0.57, Tukey’s test) (Fig. 2C).

**Figure 2.**
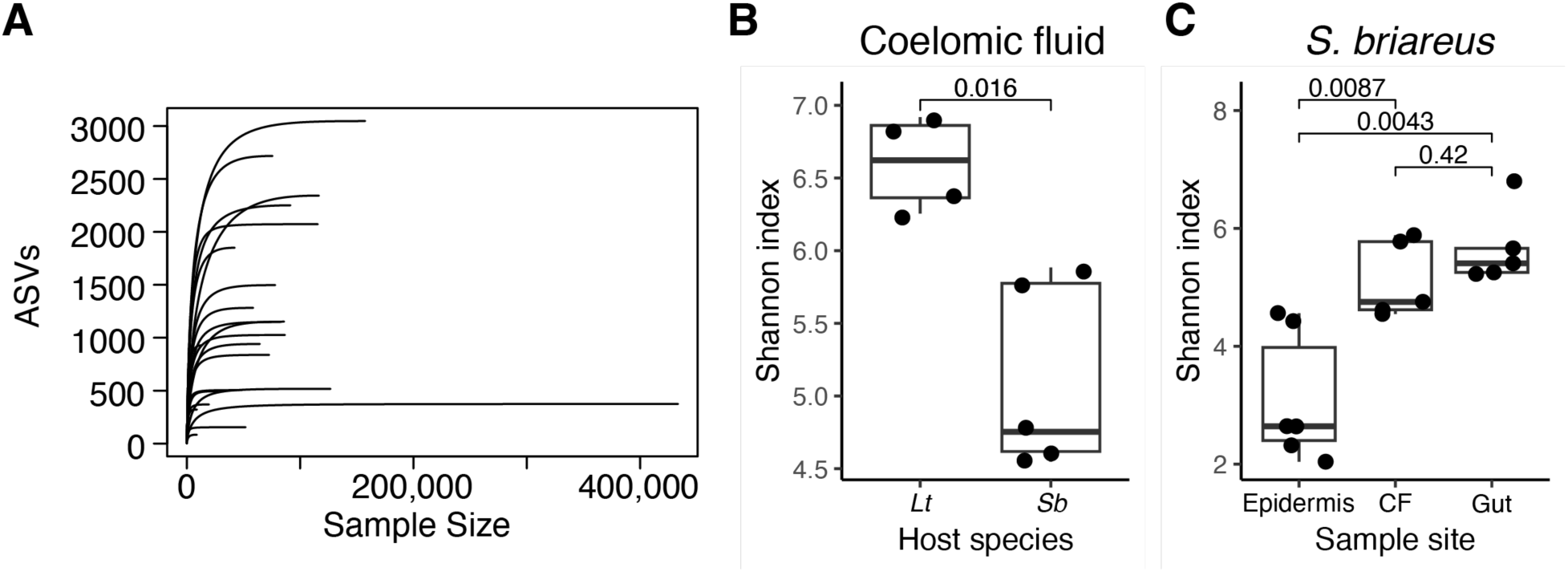
Rarefaction curves (A) and comparisons of alpha diversity (**B, C**). **A**) Rarefaction curves depict the number of ASVs recovered over iterative resampling of reads. **B, C**) Shannon diversity index of CF samples across host species (Lt = *L. tenuis*, Sb = S*. briareus*) (**B**) and across body sites among *S. briareus* samples (**C**). Lines, boxes, and whiskers indicate medians and ranges as described in Fig. 1. Values above brackets indicate p-values from a t-test (**B**) and Tukey’s test (**C**).

### Patterns in microbiome community composition

nMDS was used to assess differences in community structure across the various microbiomes sampled (Fig. 3). Sample ordination indicated differences in prokaryotic communities between Sb and Lt individuals when considering samples across all body sites (Fig. 3A), and significant differences according to host species were confirmed with PERMANOVA (F=5.4, R^2^ = 0.22, p<0.0001,). Considering both host species and body site together, host species and body site were significant drivers of community composition (PERMANOVA test: p<0.0001; F=7.5, 4.7; R^2^=0.22, 0.27; respectively;) with body site explaining slightly more variance. When analyzing only CF samples, there was again a significant difference in communities according to host species (Lt vs Sb; PERMANOVA test: F=4.2, R^2^=0.37, p=0.008). We then analyzed community variation across body sites within a single host species, Sb (Fig. 3B). Body site significantly explained a large portion of community variation (PERMANOVA test: F=5.7, R^2^=0.47, p=0.001). While there was a sizeable overlap in sample ordination by body site, there were significant differences in community composition between all three groups (PERMANOVA test: F=2.7, 5.5, 9.9; R^2^=0.25, 0.38, 0.52; p = 0.018, 0.012, 0.006; CF vs Gut, CF vs. epidermis, Gut vs. epidermis; respectively).

**Figure 3.**
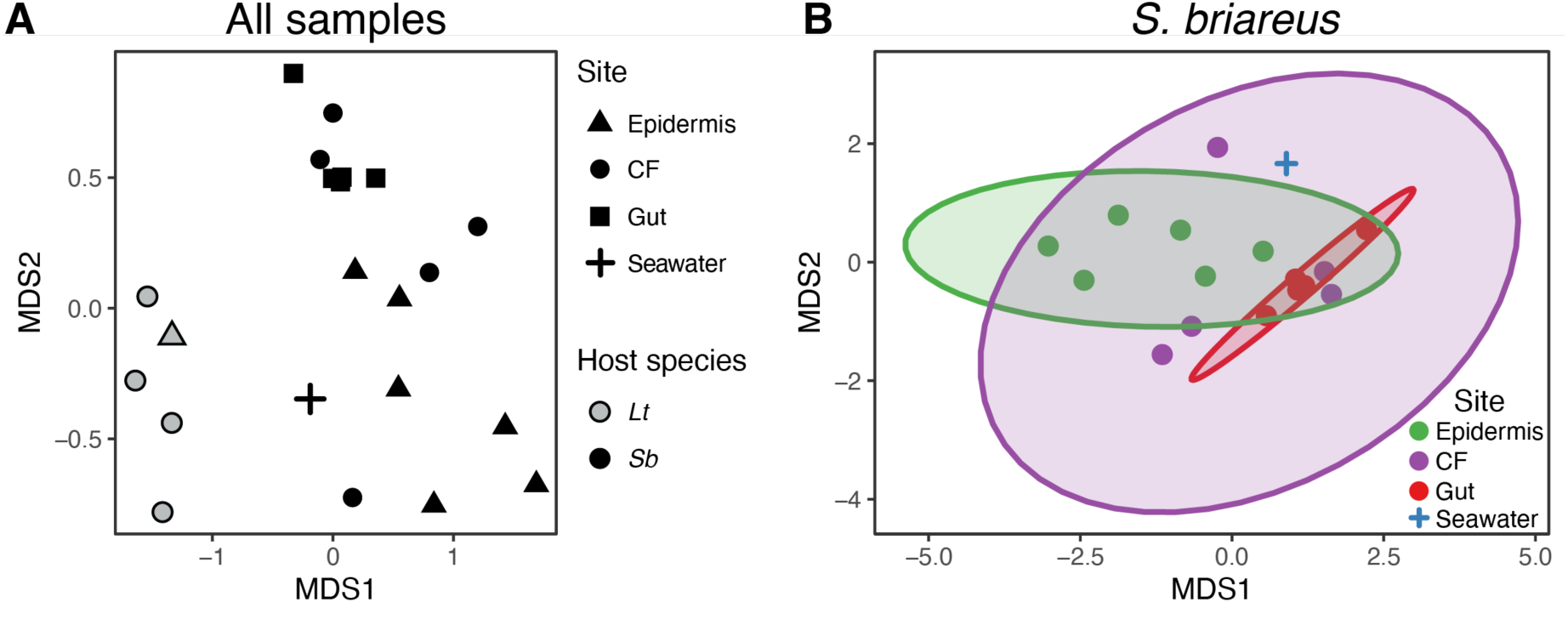
Ordination analysis of sea cucumber microbiomes. nMDS plots were constructed using nMDS of Bray-Curtis distances using ASV abundances. **A**) Analysis of all microbiome samples from *L. tenuis* (“Lt”, grey-filled symbols), *S. briareus* (“Sb”, black-filled symbols), and a seawater sample collected along with *S. briareus* samples (‘+’ symbol). Symbol shapes indicate sample type, seawater, or body site. **B**) Analysis of *S. briareus* samples and an associated seawater sample. Point shapes and color indicate sample type (body site or seawater). Ellipses depict 95% confidence intervals for the multivariate t-distribution of ordinated *S. briareus* samples. Coelomic fluid is abbreviated “CF”.

For context, prokaryotic community structure was determined for a single water sample, collected at the same time the Sb individuals were sampled. Based on nMDS ordination, this sample was somewhat distinct from the Sb samples (Fig. 3B). Without replicate water samples, however, we cannot test for any significant differences between the background water sample and Sb microbiomes. The water sample, however, did fall within the 95% confidence interval ellipse for CF samples, but not those for gut and epidermis (Fig. 3B), suggesting that surrounding seawater is rather distinct from epidermis and gut microbiomes but has some similarity to CF communities.

### Differences in specific taxa among samples

Given that significant differences were detected in whole-community analysis, SIMPER analysis was used to identify particular taxa driving these differences. Starting at the phylum level for CF samples, Bacteroidota were abundant in both species but were significantly higher in Lt individuals than in Sb (∼2.7-fold higher; Fig. 4A). Similarly, Bdellovibrionota was significantly higher in Lt (∼40-fold: 3.0% vs 0.07% on average). The converse was true for Desulfobacterota, Cyanobacteria, and Campylobacterota. Each of these phyla were at very low levels in Lt (means of 0.43%, 0.56%, and 0.54% of the community, respectively) but comprised ∼6-9% relative abundance on average and were ∼10 to 20-fold higher in Sb (Fig. 4A). Likewise, several orders of bacteria were significantly different between the CF of Lt and Sb, including Chitinophagales, Synechococcales, Campylobacterales, Desulfobulbales, Micrococcales, Cytophagales, Desulfobacterales, Chromatiales, and OM190 (Fig. 4B). A notable difference among these was ∼10% relative abundance of Chitinophagales in the CF of Lt, but its near absence in Sb. When comparing just Sb samples, eight phyla were significantly different in at least one - pairwise comparison between sites (Fig 5A). Most notable were higher abundances of Proteobacteria, Desulfobacterota, and Cyanobacteria in epidermis, gut, and CF samples, respectively. Pseudomonadales notably comprised ∼30% of epidermis samples but were at low abundance in CF and gut samples. Order Desulfobacterales in the phylum Desulfobacterota was most abundant in gut samples, consistent with the phylum results above. Photosynthetic cyanobacteria of the order Synechococcales, which made up the majority of cyanobacteria abundances, were significantly higher in CF samples (3.7% mean relative abundance) than in gut epidermis samples (means of 0.7 and 0.5%, respectively) (Fig. 5B). For comparison, Synechococcales only accounted for 0.5% of the community in the seawater sample, suggesting there is a possible enrichment of Synechococcales in the CF of Sb. However, as noted above, the statistical significance of this pattern could not be assessed.

**Figure 4.**
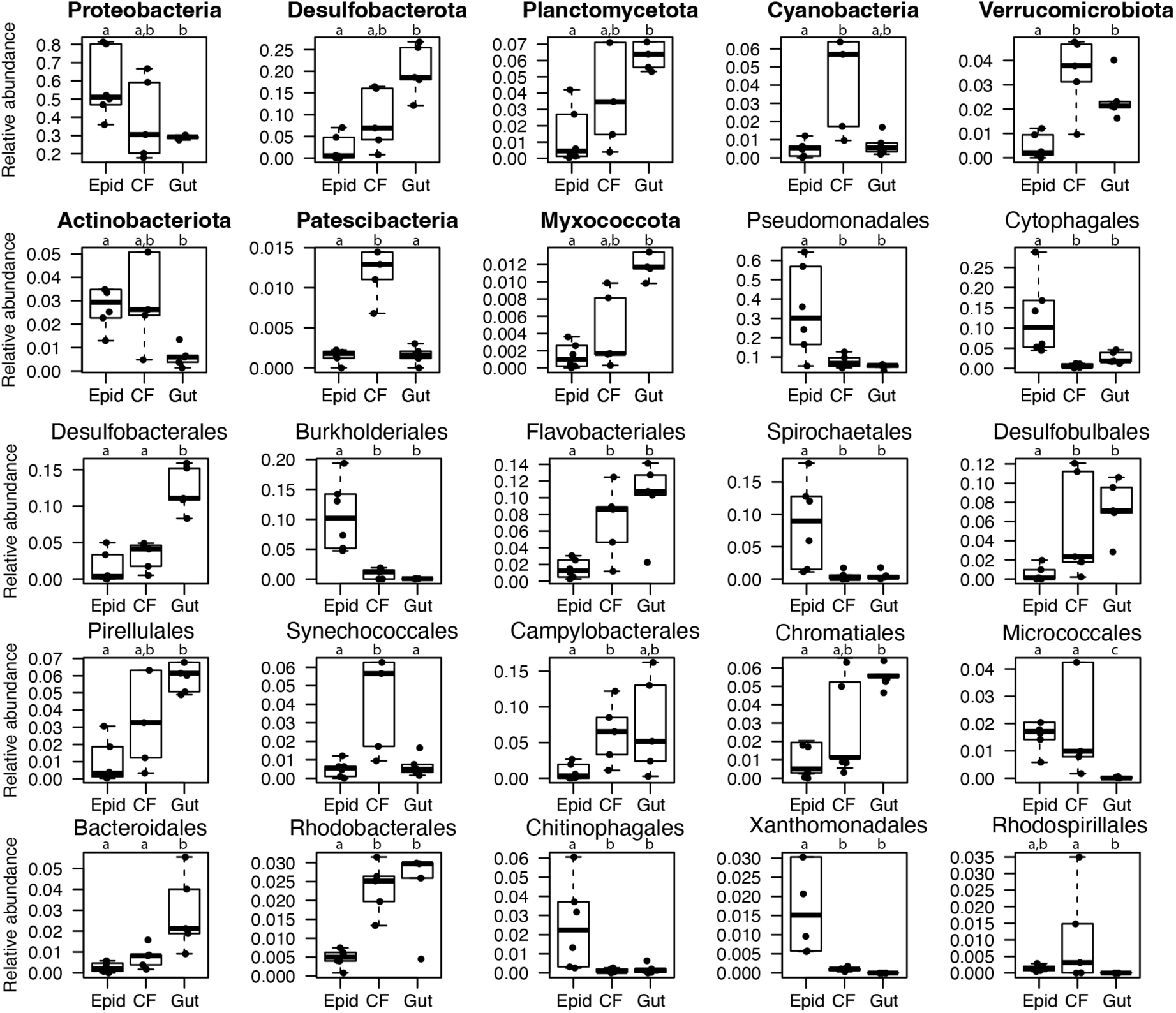
Relative abundances of phyla (**A**, phylum names in bold) and orders (**B**) that were found to be significantly different from SIMPER analysis for comparisons of CF from *L. tenuis* (Lt) vs. *S. briareus* (Sb). Lines, boxes, and whiskers depict the same information as in Fig. 1. Numbers within each panel indicate p-values from Kishino-Hasegawa tests.

**Figure 5.**
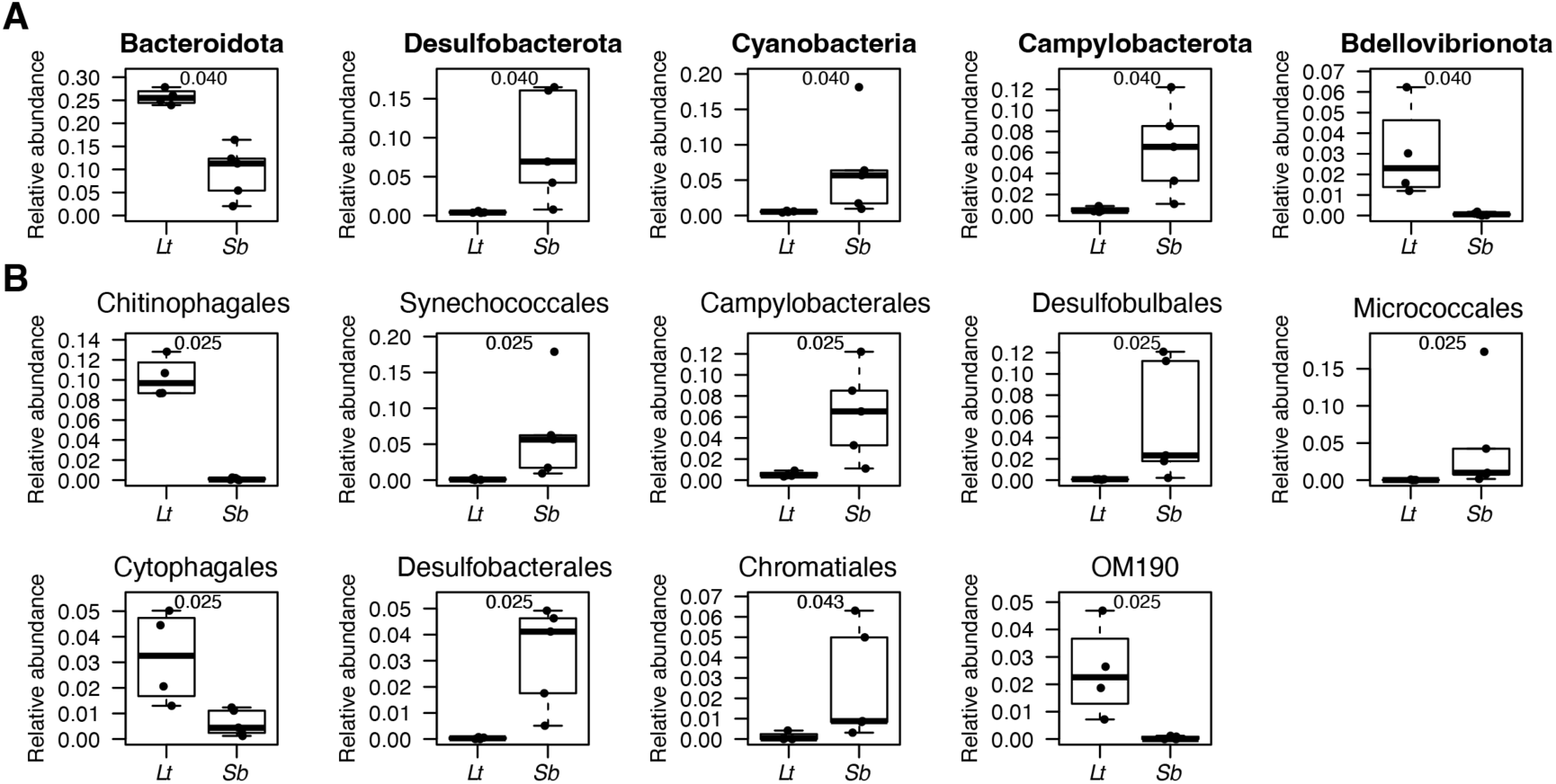
Relative abundances of phyla (names in bold in the top two rows) and orders that were found to be significantly different from SIMPER analysis for comparisons across body sites (Epid = epidermis, CF, and gut) among *S. briareus* individuals. Lines, boxes, and whiskers depict the same information as in Fig. 1. Letters above bars indicate if means were significantly different from each other, as in Fig. 1, except here with Kishino-Hasegawa tests.

## Discussion

Presented here is a focused study of the microbiomes of echinoderms found in the temperate, shallow coastal areas of the Northeast Atlantic. Only a small number of studies have examined the microbiomes of holothurians, especially in comparison to more extensive work on sea urchins and sea stars (19). This study therefore adds to the important growing knowledge of host-microbe interactions among these important marine invertebrates. The first major result is the unexpectedly high concentration of prokaryotic-sized cells in the CF of sea star *A. forbesi*, approaching 10^7^ cells/ml. Previous studies of microbial abundances in echinoderm CF are rather limited, but those report concentrations in the range of ∼10^5^-10^6^ cells/ml (19, 23, 24). We used a widely used method of DNA staining and flow cytometry to count prokaryotic-sized cells (37), while the previous studies above used staining with microscopy or qPCR. One could speculate that the high value here in the one sea star species is a technical error with our methods; however, the level of prokaryotic-sized cells in seawater was ∼10^6^ cells/ml, which is typical for surface concentrations of bacteria and archaea in seawater (19, 38, 39). Likewise, the concentrations in the CF of the three sea cucumber species were comparable to previous reports. With such high concentrations, CF may now represent a major component of the total echinoderm microbiome in *A. forbesi*. It also highlights the unusual physiology of echinoderms, which harbor high concentrations of bacteria in their internal circulatory fluids. It remains to be seen if similar high bacterial levels are found in other sea star or echinoderm species and what may drive these differences in concentrations (Fig. 1). These results warrant surveys of CF cell abundances across other echinoderm species and classes.

We found multiple differences in the microbiomes of the two sea cucumber species investigated, in terms of alpha and beta diversity. Alpha diversity patterns are not that well-studied in holothurians, but some studies have reported differences across host species and body sites in echinoderms (18, 19, 35). Such differences across hosts and body sites are common and expected for animal microbiomes, likely reflecting differences in physiology and biochemical properties of hosts and sites that in turn affects their microbiomes. Differences in community composition across sites and species are unsurprising for the same reasoning. The microbiomes of Lt and Sb were quite distinct, with host species explaining large portions of community variance (∼20-30%). Similar host- and site-associated differences in microbiome communities have specifically been documented for echinoderms (18, 19, 31, 35, 36). One limitation of this study is the lack of sequencing results across sites for Lt that would have allowed more comparisons among hosts and sites. Nonetheless, body site also explained a sizeable portion of community variance among Sb individuals (∼50%), indicating that the host species and body site have comparable strengths in shaping microbiome structure.

Specific comparisons of the CF of Lt and Sb identified several distinct patterns in specific bacterial taxa. However, many of these taxa (both phyla and orders) contain diverse members, so it is not fully clear what biological importance these patterns have. For example, Lt had higher levels of Bacteroidota. This phylum contains many diverse taxa, but many are anaerobic bacteria (40), including two orders within, Desulfobulbales and Desulfobacterota, which were both higher in the CF of Sb. This perhaps suggests that Sb harbors more sites with anaerobic conditions within its coelom than Lt. The order Chitnophagales contains chitin degraders (41), so perhaps the CF of Lt contains more chitin to support this difference. Another intriguing result is the higher prevalence of the cyanobacterial order Synechococcales in Sb CF (see below).

Comparisons among the single host, Sb, revealed interesting differences in microbiome communities. The epidermis harbored distinct communities from gut and CF samples while there was a decent overlap in the gut and CF communities (Fig. 3). This pattern is perhaps unsurprising because echinoderm digestive organs are surrounded by CF such that there is a stronger potential for exchange of microbes and perhaps more similar biochemical conditions between the two. Furthermore, the epidermis is external and a very different tissue: it excretes mucus, is exposed to seawater, and unsurprisingly supports a distinct community. As noted above, particular phyla and orders associated with these differences often contain diverse taxa without clear coherent physiologies, but we do discuss a few interesting patterns. Epidermis samples were enriched in Pseudomonadales, Burkholderiales, Spirochaetales, and Chitinophagales. The orders Pseudomonadales and Burkholderiales contain many pathogenic taxa. The former includes members of the genus *Pseudomonas*, some of which have been associated with sea star wasting disease (42). However, since these two orders contain non-pathogens, it is unclear what is the biological importance of these patterns, and all of the individuals sampled here appeared to be healthy. Spirochete taxa have been documented in a few studies in the CF and guts of echinoderms (34) (19, 43). Here we find them at higher abundance in the epidermis. Jackson et al. similarly found a higher prevalence of spirochaetes in hard tissues (gonads and body wall) compared to CF. The higher presence of Chitinophagales on epidermis sites makes sense since these taxa likely are exposed to more chitin from seawater sources, but this pattern conflicts with the observation noted above of their appreciable levels in the CF of Lt.

Several taxa displayed an apparent gradient of increasing relative abundance from epidermis to CF and gut samples. This included the phyla Proteobacteria, Desulfobacterota, Planctomyceota, and Myxococcota. The former contains the orders Desulfobacterales and Desulfobulbales, and several other orders had this gradient-like pattern (Flavobacteriales, Pirellulales, Chromatiales, Micrococcales, Bacteroidales, Rhodobacterales). These patterns could reflect increasing nutrient and anaerobic conditions from surface to internal sites, with the gut likely having the lowest oxygen levels. This is at least congruent with patterns of increasing anaerobes, i.e. Desulfobacterota, Desulfobacterales, and Desulfobulbales. Other studies, concordantly find high levels of Desulfobacterales in echinoderm gut samples (43). Higher levels of Bacteroidales in gut samples are consistent with the fact that this order contains many anaerobic taxa and the common association of the phylum Bacterioidetes with echinoderm guts (43) and many animals (44). However, again, it is hard to define coherent physiologies and ecologies for these broad taxa. In sum, there appear to be several sharp differences in bacterial taxa across hosts and body sites that provide potential avenues for further investigation of physiological differences among sea cucumbers and their tissues. Since echinoderm microbiome studies are still growing, it remains to be seen if some of these patterns hold up more generally.

Cross-site comparisons revealed curiously higher levels of cyanobacteria (order Synechococcales) in the CF of Sb (means of 6.5%, 0.51, and 0.67% for CF, epidermis and gut, respectively). Synechococcales, also occurred at a lower relative abundance in seawater (0.51%), although we cannot test the significance of this pattern. Together these results suggest that these cyanobacteria are specifically enriched in the CF of Sb. This is surprising since cyanobacteria are photosynthetic and there likely is insufficient light inside the animal to sustain photosynthetic growth. Cyanobacteria and Synechococcales members are capable of heterotrophic and/or mixotrophic growth (45). *Prochlorococcus*, the sister genus, can also re-grow after being in the dark for up to 11 days (46). These provide possible mechanisms for persistence within CF, but it is unclear if their presence is simply transient or perhaps they have some ‘active’ role in the CF microbiome.

This pattern highlights how more generally microbes in the background seawater sample differed somewhat from associated Sb CF samples. As previously stated, since only one seawater sample was analyzed, we cannot assess any significant differences between host and background seawater. It is still unclear how interactions of background seawater influence CF communities. Current understanding is that the pore-like madreporite structure in echinoderms regulates the flow of seawater into the coelom, albeit at slow rates (<5% of body weight per day) (47–49). Bacteria presumably are imported through the pores whose diameters are much larger (>20 µm) than bacteria (47, 49). Others have suggested that CF microbiomes represent transient components of seawater microbes with evidence of selection of particular taxa in comparison to seawater as we see here (18, 19, 36). How CF communities are established and maintained in relation to seawater input clearly requires further investigation.

In summary, this work adds to the growing knowledge in the study of echinoderm microbiomes, which is still nascent. Many studies have focused on species from the Pacific and/or tropical echinoderms, so this study helps expand knowledge about temperate, Atlantic host species. Our results confirm broad patterns of host-specific communities and clear differences across body sites, likely reflecting selective, deterministic processes. Given relatively low sampling across the vast diversity of echinoderms (∼7000 species), it remains to be seen what global patterns may emerge, such as the association of particular taxa with certain host body sites or the degree of overlap between CF with gut or background seawater. Large-scale microbial analysis of host taxa, for example in insects and vertebrates, has been valuable in elucidating key factors shaping microbiomes, such as locomotion, lifestyle, and diet (50–52). Applying such broad sampling of echinoderms would unlock key knowledge about their microbiomes, for example how various feeding modes shape their associated gut microbes.

## Materials and Methods

### Sample collection

Three species of echinoderms were collected on September 16^th^, 2020, by the Marine Biological Laboratory (MBL) Marine Resource Center: *Leptosynapta tenuis* (sea cucumber)*, Sclerodactyla briareus* (sea cucumber)*, and Asterias forbesi* (sea star). Animals were collected from Waquoit Bay, located off the coast of Massachusetts, using net-dragging techniques aboard the R/V Gemma. Six individuals of each species were held separately in tanks containing background seawater for 72 hours until they were transported to the lab at Clark University for processing. In the lab, 10 ml of seawater was filtered through a 0.22 μm Sterivex filter. One ml of background water was preserved with 0.125% glutaraldehyde and frozen and stored at -80 °C.

Epidermis microbiome samples were collected by swabbing a 1 cm square area from the ventral end of each animal. Swabs were stored at -80 °C for future DNA extraction. Coelomic fluid was extracted from each animal by inserting a sterile 0.21-gauge needle at a 45-degree angle into the ventral side of the organism and slowly extracting fluid. One ml of CF was preserved with 0.125% glutaraldehyde, and another ml of CF was captured by gentle filtration onto a 0.22 µM PVDF Sterivex™ filter for DNA extractions. Samples were frozen and stored at -80 °C for further DNA extraction. Echinoderms were then dissected in a disinfected dissection tray. The tray, scalpels, and tweezers were disinfected with 10% bleach and 70% ethanol in between each organism. Each organism was carefully dissected laterally to preserve the gut structure. The material from the gut was removed frozen and stored at -80 ° C.

### Flow cytometry

Prokaryotic-sized cell abundances in CF and seawater samples were measured by flow cytometry, generally following the approach of Marie et al. (37) using a Beckman Coulter Cytoflex flow cytometer. Preserved samples were thawed and stained with 1X Sybr Green I for at least 10 minutes in the dark, at room temperature before running flow cytometry analysis. Particles were excited with a 488 nm laser and side scatter light and green fluorescence were measured, the latter using a 525 ± 40 nm band pass filter. Particle data was collected by triggering on the green fluorescence channel. After each sample, the machine sample line was rinsed with 10% bleach followed bv approximately 0.2 ml of purified water to limit cross-contamination between samples. Prokaryotic-sized cell abundances (cells/ml) were determined by gating particle populations that were positively stained and in roughly a 0.5 to 2 µm size range, based on side scatter levels referenced to beads of known sizes. Positively stained particles were gated as those with green fluorescence distinct and above noise populations of negative control samples: purified water stained with Sybr Green I, as above. Each sample was run on the flow cytometer in triplicate and the average cell abundance of each sample was used for subsequent analyses.

### DNA Extraction and 16S rRNA amplification

DNA was extracted using a phenol:chloroform method (53) starting with 450 µl of lysis buffer: 0.1 M NaCl, 10mM Tris-Cl pH 8.0, 10mM EDTA pH 8.0, and 10% sodium dodecyl sulfate. Filters of CF or seawater or 0.1 g of gut material were placed in tubes with lysis buffer and vortexed briefly. For epidermis samples, the tips of swabs were cut off with cleaned scissors (wiped with 10% bleach and 70% ethanol) into tubes with lysis buffer and vortexed for 1 minute. Sample tubes with lysis buffer were then boiled in a water bath for 2 minutes. Tubes were then spun down briefly. Roughly 500 μl of lysed sample was used for serial, equal-volume extractions with phenol, 1:1 phenol:chloroform, and 25:24:1 phenol:chloroform:isoamyl alcohol (pH 8) whereby samples were mixed by inversion five times, centrifuged at 13,000 x g for two minutes, and the resulting aqueous phase removed and passaged. DNA was precipitated by adding 0.25X volume of 10.5M ammonium acetate, 2X of ice-cold 100% ethanol, and stored at -20 °C overnight. DNA was centrifuged for 90 minutes at 20,800 x g at 4 °C in a refrigerated centrifuge. The supernatant was removed, and the DNA pellet was washed in 250 µl of cold ethanol followed by centrifugation at 20,800 x g for five minutes. After the ethanol was removed and the pellet air-dried for several minutes, DNA was resuspended in 15 μl of 40 °C 10 mM Tris, 1 mM EDTA pH 8.0 at room temperature for one hour before quantification with a Qubit instrument with DNA high sensitivity kit and a Nanodrop instrument to assess purity.

The V4-V5 region of the 16S rRNA gene was amplified by PCR with primers 515Y and 806R linked to barcoded and indexed adaptors following the scheme outlined by Parada et al. (54). Twenty-five µl PCR reactions contained Q5 Hot Start High-Fidelity polymerase and buffer (New England Biolabs; Ipswitch, MA), 5 µM of primers, and 5 µl of DNA or water for negative controls. Products were visualized by agarose gel electrophoresis and purified using magnetic beads (55). Samples were quantified using a Qubit instrument as above. Products were diluted to 2 ng/µl and equal volumes of samples were pooled with 5 µl each of negative control reactions. This pooled sample was purified again with magnetic beads. The library was sequenced at the University of Connecticut Microbial Analysis, Resources, and Services (MARS) facilities on an Illumina MiSeq instrument with a v2 reagent kit.

### Microbiome analysis

Sequencing data was provided by the sequencer as demultiplexed by the reverse primer index, and samples were demultiplexed by the forward primer barcode using Sabre (56) with default settings after trimming a four base degenerate pad following the 515F primer. Forward and reverse sequencing reads were then processed in Qiime2 (57) following the pipeline outlined in Yeh et al., 2019, and outlined here in brief. Trim lengths for forward and reverse read trimming lengths were set at 221 and 229 bp, respectively after examining read quality with the ’demux summarize’ function. These were the longest possible reads lengths for which 91% of reads still had mean Phred quality scores (Q) of >30. DADA2 (58) was used to merge reads and generate amplicon sequence variances (ASVs) using default settings and the trim lengths above. ASVs were classified using the SILVA 132 database (59) with the “feature-classifier classify-learn” function. Only echinoderm and seawater samples with greater than 8,000 reads were included in subsequent analyses. For the 22 samples that passed this requirement, 7,000 reads each was randomly resampled 20 times and read counts were averaged to generate equal read depths for subsequent analyses.

Statistical analyses were done in R Studio and most microbial community analysis was done with functions in the vegan package (60). T-tests, Analysis of Variance (ANOVA), and Tukey tests were done with base R functions *t.test*, *aov*, and *TukeyHSD* respectively. ASV richness and Simpson diversity index were computed with the function *diversity*. Rarefaction curves were generated with the function *rarecurve*. Non-metric multidimensional scaling plots were generated with Bray-Curtis dissimilarity scores with the function *metaMDS*, and permutational multivariate analysis of variance (PERMANOVA) was done using the function *adonis2*. Similarity percentage analysis (SIMPER) was done to identify particular phyla and orders with mean abundances of at least 1% that contributed significantly to differences in microbial communities using the function *simper* and wrapper scripts simper_pretty.R and R_krusk.r (61) to iterate over taxa and perform post-hoc Kruskal-Wallis tests with Benjamini and Hochberg correction.

## Data availability

Sequence read data are available at the National Center for Biotechnology Information under BioProject accession PRJNA1281654.

## Acknowledgements

We thank the staff at Marine Biological Laboratory Marine Resource Center who collected the animals used in this study. The authors declare no conflict of interest. This work was supported in part by funds from the Clark University Biology Department.

## Notes

### Competing Interest Statement

The authors have declared no competing interest.

